# Insertion of an endogenous Jaagsiekte Sheep Retrovirus element into the *BCO2 -* gene abolishes its function and leads to yellow discoloration of adipose tissue in Norwegian Spælsau (*Ovis aries*)

**DOI:** 10.1101/2020.06.11.145755

**Authors:** Matthew Kent, Michel Moser, Inger Anne Boman, Kristine Lindtveit, Mariann Árnyasi, Kristil Kindem Sundsaasen, Dag Inge Våge

## Abstract

**Background:** The accumulation of carotenoids in adipose tissue leading to yellow fat is, in sheep, a heritable recessive trait that can be attributed to a nonsense mutation in the *beta-carotene oxygenase 2 (BCO2)* gene. However, not all sheep breeds suffering from yellow fat have this nonsense mutation, meaning that other functional mechanisms must exist. We investigated one such breed, the Norwegian spælsau.

**Results:** In spælsau we detected an aberration in *BCO2* mRNA. Nanopore sequencing of genomic DNA revealed the insertion of a 7.9 kb endogenous Jaagsiekte Sheep Retrovirus (enJSRV) sequence in the first intron of the *BCO2* gene. Close examination of its cDNA revealed that the *BCO2* genes first exon was spliced together with enJSRV-sequence immediately downstream of a potential - AG splice acceptor site at enJSRV position 415. The hybrid protein product consists of 29 amino acids coded by the *BCO2* exon 1, followed by 29 amino acids arbitrary coded for by the enJSRV-sequence, before a translation stop codon is reached.

**Conclusions:** Considering that the functional BCO2 protein consists of 575 amino acids, it is unlikely that the 58 amino acid BCO2/enJSRV hybrid protein can display any enzymatic function. The existence of this novel *BCO2* allele represents an alternative functional mechanism accounting for *BCO2* inactivation and is a perfect example of the potential benefits for searching for structural variants using long-read sequencing data.

## Background

The yellow coloration of adipose tissue in sheep is known to be a heritable trait in different sheep breeds and is caused by the accumulation of carotenoids [1–4]. In terms of consumer preferences, “yellow fat” is an undesirable meat quality that leads to loss of product value and is therefore not wanted in meat animals.

In animals, two enzymes displaying different cleavage effects on carotenoids have been identified; *BCO1* (*beta-carotene oxygenase 1*) which cleaves β-carotene symmetrically into two retinal molecules, and *BCO2* (*beta-carotene oxygenase 2*) which cleaves β-carotene asymmetrically into β-apo-10’-carotenal (C27) and β-ionone (C13) [5–8].

Mutations influencing either transcript levels or protein coding sequence have been reported for both genes in several organisms. Mutations in *BCO2* especially seem to result in accumulation of carotenoids [9, 10], possibly due to its broader substrate specificity [11, 12].

Earlier we have reported that a nonsense mutation in the coding sequence of ovine *BCO2* is associated with yellow fat in Norwegian White Sheep breed [13]. However, in the native Norwegian breed Spælsau, the yellow fat trait is found to segregate in the absence of this mutation, suggesting that alternative functional mechanism is influencing this trait. In the present study we searched for novel *BCO2* mutations in the Spælsau breed that could explain the yellow fat phenotype.

## Results and discussion

Our initial pilot experiments to better understand the mechanism(s) responsible for yellow fat included performing PCR on cDNA from animals presenting normal or yellow fat to detect expression of *BCO2*. Four primer pairs targeting *BCO2* produced 4 fragments of expected size in the three individuals showing normal fat, while no fragments were amplified from the individual with yellow fat (Figure 1). Our provisional conclusion from this was that the yellow fat phenotype was due to downregulated expression of *BCO2* mRNA, but our results did not offer insight into the mechanism underlying this.

**Figure 1.**
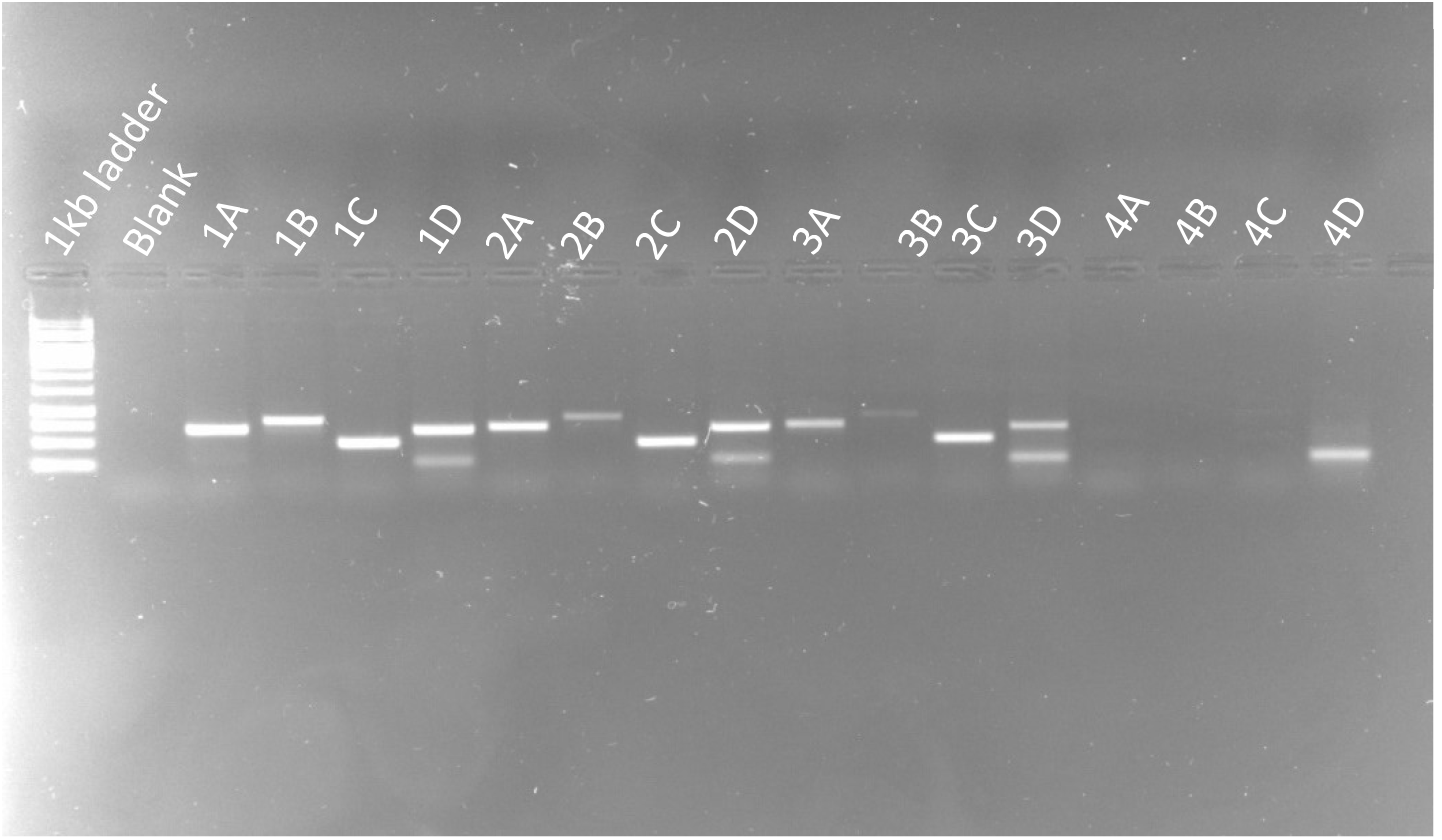
Figure is showing 4 different sets of primers (A: 7541-7542, B: 7543-7544, C: 7545-7546, D: 7547-7548) amplifying different subregions of the ovine BCO2 cDNA sequence in 4 different individuals (20025, 50289, 70203 and 70346, respectively). No fragments are amplified in the yellow fat individual (70346), the band in 4D is an artefact which also is visible in the 3 other individuals (in addition to the band of expected size found in the other 3).

To investigate if novel genomic rearrangements in the *BCO2* region could account for the lack of detectable mRNA, genomic DNA from the single yellow fat individual (70346) and from an individual with the normal fat phenotype (20025) were sequenced using nanopore long read technology. Over the course of two consecutive PromethION sequencing runs, one flow-cell yielded 6,908,462 reads (73 Gb raw data). After demultiplexing, 78% of reads (n = 5,387,322) were assigned either to sample 20025 (1,330,100 reads; 21 Gb) or to sample 70346 (2,512,544 reads; 27.2 Gb). After filtering for quality and read length (Q>7, length > 4 kb) a total of 20.2 Gb data remained for individual 20025 (≈7X coverage; median read length = 14,792, N50 read length = 23,141) and 25.8 Gb for individual 70346 (≈9X coverage, median read length = 10,567, N50 read length = 13,570 bp).

Filtered nanopore reads were mapped to the Oar_rambouillet_v1.0 reference and examined for SV’s. Within the 70kb genomic interval harbouring *BCO2* (NC_040266.1: 25,021,687-25,091,194), 5 SV’s were detected. Four of these were found in both individuals (normal and yellow fat) and may represent a spælsau breed-specific difference. These were disregarded as candidate SVs explaining the yellow fat phenotype. The single remaining SV consisted of a 7,938 bp insertion (Figure 2) at reference genome position NC_040266.1: 25,022,547 which is 730 bp downstream from the end of *BCO2*’s first exon. The inserted sequence showed high similarity (99.04 % identity) to an endogenous Jaagsiekte sheep retrovirus (enJSRV; accession MF175071.1). The enJSRV insertion was verified with PCR using primer pairs 7554-7555 and 7556-7557 in the two sequenced animals (20025 and 70346), which gave products (of expected size) spanning the upstream and downstream sheep-virus junctions, respectively. Under the PCR conditions we used, the primer pair spanning the insertions site 7552-7553 did not amplify any product in the yellow fat animal (70346), while it did in the heterozygous normal animal (20225).

**Figure 2.**
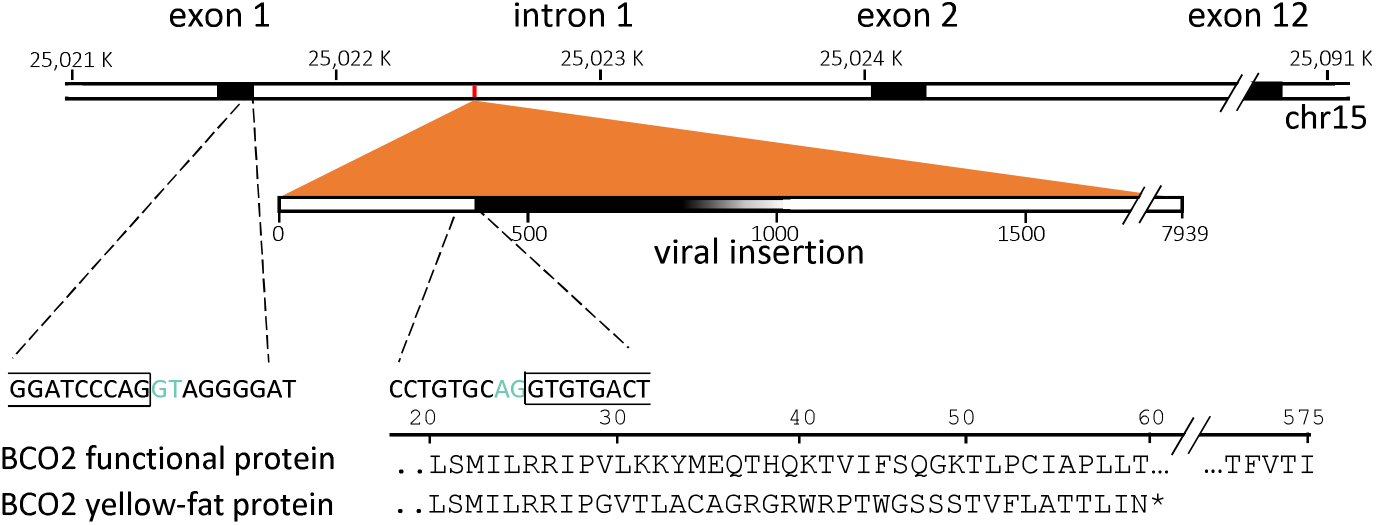
Diagram of the viral insertion in the *BCO2* gene (NC_040266.1: 25021687-25091194). The splice donor and acceptor sites are indicated in green.

After establishing that an enJSRV-sequence had been inserted between exon 1 and exon 2 in the *BCO2* gene, we tested whether exon 1 (located upstream of the insertion point) could still be expressed in a homozygous carrier by amplifying a subregion of exon 1 (primers 7550-7551) from cDNA. A band of expected size (112 bp) was produced, indicating that *BCO2* exon 1 was being transcribed in individual 70346. However, since using PCR primers located downstream of the insertion point did not yield any products from cDNA (see Figure 1), the next step was to test the hypothesis that the enJSRV – sequence within intron 1 could have interrupted the normal splicing process.

By combining a *BCO2* exon 1 forward primer (7550) with enJSRV specific reverse primer (7555) we were able to generate a 188 bp PCR product using cDNA from the yellow fat individual. Sanger sequencing this fragment between the two primer binding sites revealed it to be composed of 94 bp from *BCO2* exon 1, and 55 bp of enJSRV sequence, beginning at the base corresponding to position 415 in the MF175071.1 sequence (Figure 2). If translated, this hybrid mRNA sequence would give rise to 29 amino acids of the wild-type ovine BCO2 protein chimerized to 1 aa coded by the junction sequence followed by 28 amino acids coded by the enJSRV-sequence before the protein is terminated by a stop codon. Knowing that the full-length protein consists of 575 amino acids, we theorize that it is highly unlikely that these 58 peptides can perform the functions of wildtype BCO2 enzyme in animals homozygous for the enJSRV insertion.

These preliminary observations suggesting a functional relationship between the enJSRV insertion and *BCO2* activity were based upon a very small number of individuals. Our analysis of sample set 2, which consisted of 26 individuals with an expected relatively high frequency of the yellow fat allele, showed that 10 were heterozygous for the insertion, 4 were homozygous for the insertion, and the remaining 12 lambs were wild-type. In the abattoir-report the 4 homozygous animals were reported to have yellow fat, while the remaining group had white fat (Table 2).

## Conclusions

In this study we identified an insertion of an endogenous Jaagsiekte Sheep Retrovirus element in the *BCO2* gene in the Norwegian spælsau breed by using Nanopore sequencing technology. The insertion is localised in the first intron of *BCO2* and interrupts the mRNA splicing process. The resulting mRNA consists of *BCO2* exon 1 sequence fused with endogenous retrovirus sequence, leaving an open reading frame of totally 58 amino acids. Given that the wild type BCO2 enzyme consists of 575 amino acids, we find it highly unlikely that the hybrid protein of 58 aa can have any enzyme function. Our results make it possible for the sheep farming industry to screen their breeding candidates for this variant to avoid the yellow fat phenotype. We also think this paper clearly illustrate the strength of using long-read sequencing technology to search for structural variation and it also exemplifies how such variation can directly influence gene function.

## Material and methods

### Animals

In Norwegian abattoirs, fat colour in sheep carcasses is graded as “normal” or “yellow” and recorded at the individual level within the national Sheep Recording System. To perform a pilot analysis, we obtained samples from 4 Spælsau animals at slaughter from a farm with a history of producing yellow fat lambs (sample set 1). Liver samples (approximately 0.125 cm^3^) from a “yellow” male (individual 70346), and from two “normal” males (50289,70203) and one “normal” female (20025) were collected and stored in RNAlater™ (QIAGEN, Hilden, Germany). The normal individuals may be carriers of the genetic variant(s) causing yellow fat, based on their relationship to known “yellow” animals, but most likely not homozygous.

Subsequently, a larger set of 26 Spælsau lamb samples were collected from two farms in Norway (sample set 2). Lambs from each farm were half-sibs descending from one of two rams registered with some offspring displaying the yellow fat trait. Similarly, mothers of these lambs were also known to be related to individuals presenting yellow fat, and therefore had an increased probability of carrying at least one yellow fat allele.

### RNA extraction and cDNA synthesis

RNA was extracted using the RNeasy®Plus Universal Mini Kit from QIAGEN according to the manufacturer’s instructions. The concentration and purity of the RNA was measured using a NanoDrop 8000 (Thermo Scientific), and the integrity of RNA was measured using a Bioanalyzer 2100 (Agilent). All samples had an RNA integrity value (RIN) of at least 8.1. cDNA was produced using a SuperScript™ II Reverse Transcriptase kit from Invitrogen according to manufacturer’s instructions.

### PCR amplification of BCO2 cDNA

To amplify the complete cDNA sequence of *BCO2*, four pairs of validated primers (Våge & Boman 2010) were mixed with cDNA and AmpliTaq Gold® Polymerase (Applied Biosystems) in four separate PCR reactions generating overlapping regions of the *BCO2* cDNA [13]. The PCR was performed using 10 min at 95°C, and 40 cycles of 95°C for 30 sec, 57°C for 30 sec and 72°C for 1.5 min.

### Nanopore sequencing

High-molecular-weight DNA from white (20025) and yellow fat (70346) individuals was extracted from liver tissue using Genomic-tips (G/100) from Qiagen, and fragments smaller than 25Kb were progressively depleted using a Short Read Eliminator kit (Circulomics; USA). Sequencing libraries were prepared using Ligation sequencing kit (LSK109, Oxford Nanopore) with native barcoding (EXP-NBD104, Oxford Nanopore). Equal masses of libraries were combined, 300 ng was loaded into a single PromethION flowcell (FLO-PRO002). After 23 hours the run was stopped (producing 36 Gb raw data) and the flow-cell regenerated according to Oxford Nanopore’s nuclease flush protocol. The remaining pooled library (230 ng) was added to the same flow-cell and a new sequencing run performed (22.2 hours; 22 Gb raw data).

### Base-calling and quality filtering

Nanopore reads were base-called using Guppy, version 3.0.3 (Oxford Nanopore, MinKNOW v.19.05.1) using the ‘High accuracy’ flip-flop model. Reads from the two sequencing runs were merged and demultiplexed with QCAT (https://github.com/nanoporetech/qcat).

Demultiplexed reads from each sample were trimmed for 50 bp from the start and quality-filtered for average PHRED quality > 7 and minimum length of 4000 bp with fastp (v0.19.5) options “ – disable trim_poly_g --disable_adapter_trimming -q 7 -l 4000 -f 50” [14].

### Structural variant detection

Quality-filtered reads were mapped to the sheep reference genome (RefSeq accession GCF_002742125.1) using minimap2 (v2.16-r922) [15]. Structural variants (SV’s) were called with the SV-caller SVIM (v0.5.0) using default parameters [16]. SVs within 100 Kbp of the candidate gene *BCO2* (NM_001159278.1) were inspected manually.

As a complementary approach to identify larger rearrangements between the sheep reference genome and the yellow fat individual, quality-filtered reads of sheep_70346 were assembled using Flye software (v2.4.2) with options “--genome-size 2.8g -m 10000” [17]. Contigs from the sheep_70346 genome assembly were mapped to the sheep reference genome using minimap2 (v2.16-r922) [15]. Candidate contigs spanning *BCO2*-region were aligned and dot-plotted against the sheep reference genome using Gepard (v1.40) for visual inspection [18].

### Identification of intronic insertion in BCO2

Candidate SVs within the *BCO2* gene region were filtered for presence only in sheep_70346 or following expected allele pattern among the two individuals (homozygous in sheep_70346 and heterozygous in sheep_20025). On-line blastn [19] was used to identify the inserted sequence in *BCO2* intronic region (INS_chr15_25022547). Insertion boundaries were determined by aligning contigs of sheep_70346 genome assembly against the Oar_rambouillet_v1.0 reference genome.

### Detection of BCO2 exon 1 and BCO2-JSRV hybrid cDNA in the yellow fat animal

A 112 bp exon 1 fragment of *BCO2* from the yellow fat individual was amplified using primers 7550/7551 (Table 1). A hybrid fragment consisting of sheep *BCO2* exon 1 sequence and JSRV sequence were amplified from the same individual using primers 7550/7555. In both cases cDNA was used as template with the following thermo cycler program: 10 min at 95°C and 40 cycles at 95°C for 30 sec, 57°C for 30 sec and 72°C for 30 sec using AmpliTaq Gold® Polymerase (Applied Biosystems). The 7550-7555 product was DNA sequenced by Sanger technology at Eurofins Scientific, Luxembourg, using the same two primers for the sequencing reaction.

**Table 1.**
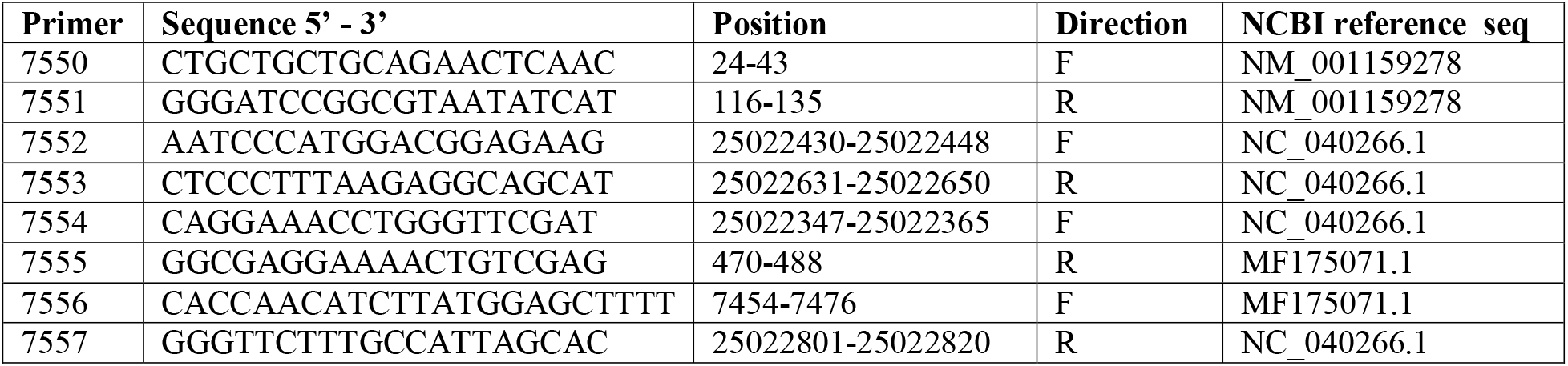
Primers used in addition to those described in Våge & Boman 2010.

**Table 2.** Fat colour grading and genotypes of 26 lambs descending from two different rams (76245736 and 89012427) known to produce yellow fat offspring. The ewes also have an increased probability of carrying the yellow fat allele, due to relationship to known carriers. Genotypes are scored as wild type (wt) or carrier of the endogenous Jaagsiekte Sheep Retrovirus (ins) based on the PCR-test described in materials and methods.

| **Lamb ID** | **Ram** | **Ewe** | **Sex** | **Phenotype** | **Genotype** |
| --- | --- | --- | --- | --- | --- |
| 90027 | 76245736 | 37200184 | male | white fat | wt/ins |
| 90029 | 76245736 | 37200184 | male | white fat | wt/wt |
| 90041 | 76245736 | 89740216 | male | white fat | wt/ins |
| 90059 | 76245736 | 37198929 | female | white fat | wt/ins |
| 90060 | 76245736 | 37198929 | male | white fat | wt/wt |
| 90072 | 76245736 | 89740213 | female | white fat | wt/ins |
| 90073 | 76245736 | 89740213 | female | white fat | wt/wt |
| 90001 | 89012427 | 82424293 | female | **yellow fat** | **ins/ins** |
| 90002 | 89012427 | 82424293 | female | **yellow fat** | **ins/ins** |
| 90015 | 89012427 | 76245888 | female | white fat | wt/ins |
| 90019 | 89012427 | 88844575 | female | **yellow fat** | **ins/ins** |
| 90075 | 89012427 | 76382061 | female | white fat | wt/ins |
| 90077 | 89012427 | 76382061 | female | white fat | wt/wt |
| 90083 | 89012427 | 37199669 | female | white fat | wt/wt |
| 90104 | 89012427 | 70456589 | female | white fat | wt/ins |
| 90115 | 89012427 | 82991541 | female | white fat | wt/wt |
| 90201 | 89012427 | 88842546 | male | white fat | wt/wt |
| 90202 | 89012427 | 88842546 | male | white fat | wt/wt |
| 90214 | 89012427 | 76245888 | male | white fat | wt/wt |
| 90228 | 89012427 | 76245885 | male | white fat | wt/ins |
| 90230 | 89012427 | 76245885 | male | white fat | wt/wt |
| 90297 | 89012427 | 89400436 | male | **yellow fat** | **ins/ins** |
| 90298 | 89012427 | 89400436 | male | white fat | wt/wt |
| 90307 | 89012427 | 37199669 | male | white fat | wt/ins |
| 90330 | 89012427 | 70456589 | male | white fat | wt/wt |
| 90336 | 89012427 | 82991541 | male | white fat | wt/ins |

### PCR to detect the presence or absence of the endogenous virus sequence in genomic DNA

To detect the presence or absence of endogenous virus sequence in genomic DNA, three pairs of primers were designed (Table 1): one pair (7552/7553) was designed to amplify across the insertion point without any insertion, one pair (7554/7555) to amplify the junction between the *BCO2* intron sequence and the upstream virus region and one (7556/7557) to amplify the junction between the *BCO2* intron sequence and the downstream virus region. Expected fragments lengths are 221 bp, 684 bp and 768 bp, respectively. DNA was denatured for 10 min at 95°C and PCR run for 40 cycles at 95°C for 30 sec, 58°C for 30 sec and 72°C for 1.5 min using using AmpliTaq Gold® Polymerase (Applied Biosystems).

## Abbrevations

*BCO1*: *beta-carotene oxygenase 1*
*BCO2*: *beta-carotene oxygenase 1*
enJSRV: endogenous Jaagsiekte Sheep Retrovirus

## Declarations

### Ethics approval and consent to participate

The animals described herein are not to be considered as experimental animals as defined in EU directive 2010/63 and in the Regulation of Animal Experiments in Norway. Consequently, we did not seek ethical review or approval of this study regarding the use of experimental animals. Tissue samples used for mRNA isolation were collected from ordinary production animals in a Norwegian abattoir after the animals were dead (sample set 1). Tissue samples used for DNA isolation in the two half-sib groups were collected as part of the routine ear-tagging used on commercial farms in Norway (sample set 2).

### Consent for publication

Not applicable.

### Availability of data and materials

Raw long-range sequencing data have been submitted to the European Nucleotide Archive under accession number ERR3522460 and ERR3522461. The hybrid mRNA sequence is available under accession number MT024238. Other data are available upon reasonable requests.

### Authors’ contributions

IAB conceived the study together with DIV and MPK, identified suitable animals to sample and coordinated the sample collection. KL purified RNA/DNA and did the PCR-based experiments. MPK designed the Nanopore sequencing experiment and contributed in the overall planning of the project. MA did the Nanopore sequencing together with MPK. MM analysed the Nanopore sequencing data. KSS planned and coordinated the labwork. DIV assembled the results and made a first draft of the paper. All authors discussed and interpreted results and contributed to writing the paper. All authors have read and approved the final manuscript.

## Acknowledgements

We are grateful to the sheep breeder and the personnel at the Nortura abattoir that made this study possible. We also wish to thank Øystein W Milvang for genotyping sample set 2.

## References

1. Hill F. Xanthophyll pigmentation in sheep fat. Nature. 1962;194(4831):865–6.

2. Crane B, Clare NT. Nature of carotenoid pigments in yellow fat of sheep. N Z J Agric Res. 1975;18(3):273–5.

3. Kirton AH, Crane B, Paterson DJ, Clare NT. Yellow fat in lambs caused by carotenoid pigmentation. N Z J Agric Res. 1975;18(3):267–72.

4. Baker RL, Steine T, Vabeno AW, Breines D. The inheritance and incidence of yellow fat in Norwegian sheep. Acta Agriculturae Scandinavica. 1985;35(4):389–97.

5. von Lintig J, Vogt K. Filling the gap in vitamin A research - Molecular identification of an enzyme cleaving beta-carotene to retinal. Journal of Biological Chemistry. 2000;275(16):11915–20.

6. Wyss A, Wirtz G, Woggon WD, Brugger R, Wyss M, Friedlein A, Bachmann H, Hunziker W. Cloning and expression of beta,beta-carotene 15,15’-dioxygenase. Biochemical and Biophysical Research Communications. 2000;271(2):334–6.

7. Redmond TM, Gentleman S, Duncan T, Yu S, Wiggert B, Gantt E, Cunningham FX. Identification, expression, and substrate specificity of a mammalian beta-carotene 15,15’-dioxygenase. Journal of Biological Chemistry. 2001;276(9):6560–5.

8. Kiefer C, Hessel S, Lampert JM, Vogt K, Lederer MO, Breithaupt DE, von Lintig J. Identification and characterization of a mammalian enzyme catalyzing the asymmetric oxidative cleavage of provitamin A. Journal of Biological Chemistry. 2001;276(17):14110–6.

9. Berry SD, Davis SR, Beattie EM, Thomas NL, Burrett AK, Ward HE et al. Mutation in Bovine Beta-Carotene Oxygenase 2 Affects Milk Color. Genetics. 2009;182:923–6.

10. Eriksson J, Larson G, Gunnarsson U, Bed’hom B, Tixier-Boichard M, Stromstedt L et al. Identification of the Yellow skin gene reveals a hybrid origin of the domestic chicken. Plos Genetics. 2008;4(2).

11. Amengual J, Lobo GP, Golczak M, Li HN, Klimova T, Hoppel CL, Wyss A, Palczewski K, von Lintig J. A mitochondrial enzyme degrades carotenoids and protects against oxidative stress. FASEB J. 2011;25(3):948–59.

12. Mein JR, Dolnikowski GG, Ernst H, Russell RM, Wang XD. Enzymatic formation of apo-carotenoids from the xanthophyll carotenoids lutein, zeaxanthin and beta-cryptoxanthin by ferret carotene-9’,10’-monooxygenase. Arch Biochem Biophys. 2011;506(1):109–21.

13. Våge DI, Boman IA. A nonsense mutation in the beta-carotene oxygenase 2 (BCO2) gene is tightly associated with accumulation of carotenoids in adipose tissue in sheep (Ovis aries). BMC Genet. 2010;11:10.

14. Chen S, Zhou Y, Chen Y, Gu J. fastp: an ultra-fast all-in-one FASTQ preprocessor. Bioinformatics. 2018;34(17):i884–i90.

15. Li H. Minimap2: pairwise alignment for nucleotide sequences. Bioinformatics. 2018;34(18):3094–100.

16. Heller D, Vingron M. SVIM: Structural Variant Identification using Mapped Long Reads. Bioinformatics. 2019.

17. Kolmogorov M, Yuan J, Lin Y, Pevzner PA. Assembly of long, error-prone reads using repeat graphs. Nat Biotechnol. 2019;37(5):540–6.

18. Krumsiek J, Arnold R, Rattei T. Gepard: a rapid and sensitive tool for creating dotplots on genome scale. Bioinformatics. 2007;23(8):1026–8.

19. Coordinators NR. Database resources of the National Center for Biotechnology Information. Nucleic Acids Res. 2013;41(Database issue):D8–D20.

